# Parkinson’s disease microglia induce endogenous α-synuclein pathology in patient-specific midbrain organoids

**DOI:** 10.1101/2025.07.01.662685

**Authors:** Elisa Zuccoli, Henry Kurniawan, Isabel Rosety, Alise Zagare, Sonia Sabate-Soler, Anna-Sophie Zimmermann, Jens C. Schwamborn

## Abstract

The accumulation of misfolded α-synuclein and the loss of dopaminergic neurons are hallmarks of Parkinson’s disease (PD), contributing to the development of synucleinopathies. Although considerable progress has been made in understanding α-synuclein’s role in PD pathology, the precise mechanisms involved remain unclear. Human midbrain organoids (hMOs) have emerged as valuable models for studying PD, yet the lack of microglia limits the ability to investigate neuroimmune interactions. Recent studies show that integrating microglia into hMOs enhances neuronal maturation and functionality. Here, we generated a human midbrain assembloid model by incorporating iPSC-derived microglia into midbrain organoids from healthy control individuals and a PD patient carrying the SNCA triplication (3xSNCA) mutation. Our results show that 3xSNCA microglia alone are sufficient to induce early, endogenous formation of phosphorylated α-synuclein (pS129) pathology in the absence of exogenous fibril seeding. This PD-pathology emerged as early as day 50 of culture and was not observed in models lacking microglia. These findings highlight a critical role for patient-derived microglia in driving α- synuclein pathology and provide a physiologically relevant platform for studying early neuroimmune mechanisms in PD and testing potential therapeutic strategies.

## Introduction

Parkinson’s disease (PD) is a progressive neurodegenerative disorder characterized by the selective loss of dopaminergic neurons in the substantia nigra *pars compacta* of the midbrain and the pathological accumulation of the misfolded α-synuclein protein (Kalia and Lang, 2015; Poewe et al., 2017). A defining hallmark of PD is the formation of intracellular inclusions known as Lewy bodies and Lewy neurites, which are primarily composed of aggregated and phosphorylated α-synuclein (Spillantini et al., 1998; Goedert, 2015). These inclusions are central to the diagnosis and classification of PD and related synucleinopathies (Lashuel et al., 2013; Fujiwara et al., 2022). While the precise physiological function of α-synuclein remains under investigation, its pathological forms disrupt synaptic vesicle trafficking (Bernal-Conde et al., 2020), impair mitochondrial function (Plotegher and Duchen, 2017), and contribute to neuroinflammation and lysosomal dysfunction (Calabresi, Mechelli, et al., 2023), ultimately driving neuronal death (Stefanis, 2012; Calabresi, Di Lazzaro, et al., 2023). Furthermore, the prion-like propagation of misfolded α-synuclein between cells is thought to underlie the stereotypical progression of pathology observed in PD (Lashuel et al., 2013), with phosphorylation at Ser129 exacerbating mitochondrial dysfunction and oxidative stress (Kawahata et al., 2022).

Genetic alterations in the *SNCA* gene—including point mutations, duplications, and triplications—cause autosomal dominant forms of PD and are strongly associated with increased α-synuclein expression and the development of Lewy pathology (Polymeropoulos et al., 1997; Krüger et al., 1998; Zarranz et al., 2004; Ibáñez et al., 2004; Singleton et al., 2003). Despite major progress in understanding α-synuclein’s role in disease, the mechanisms underlying its pathological accumulation and downstream effects remain unclear (Sulzer & Edwards, 2019; Burré, 2015).

Human midbrain organoids (hMOs), derived from induced pluripotent stem cells (iPSCs) (Takahashi & Yamanaka, 2006), have emerged as powerful *in vitro* systems for modelling PD. These self-organizing, three-dimensional cultures recapitulate essential features of the human midbrain, including the presence of mature dopaminergic neurons, astrocytes, and oligodendrocytes, along with dopamine release and electrophysiological activity (Monzel et al., 2017). hMOs have been successfully used to model several genetic forms of PD, including mutations in LRRK2, GBA, or PINK1, as well as the triplication in the *SNCA* gene (Smits et al., 2019; Kim et al., 2019; Rosety et al., 2023; Jarazo et al., 2022; Muwanigwa et al., 2024). However, a key limitation of standard hMO systems is the absence of microglia - the brain’s resident immune cells - due to their mesodermal origin.

Microglia, which represent 5% to 15% of the brain cells in adults, are tissue-resident macrophages derived from yolk sac progenitors during early embryogenesis (Thion et al., 2018; Ginhoux et al., 2010; Li & Barres, 2018; Schulz et al., 2012). Their omission from hMOs restricts the ability to model critical immune-mediated mechanisms of neurodegeneration, such as synaptic pruning, phagocytosis of apoptotic neurons, and neuroinflammatory responses, all of which are central to Parkinson’s disease (Tremblay et al., 2011; Wake et al., 2009; Ormel et al., 2018). Microglia play a key role in maintaining brain homeostasis and immune defence, interacting with neurons, astrocytes, and oligodendrocytes, and their dysfunction in PD is closely associated with chronic neuroinflammation (Shabab et al., 2017). Sabate-Soler et al. (2022) demonstrated that incorporating microglia into hMO systems leads to enhanced neuronal maturation and functionality, providing a more comprehensive model of the human midbrain and its cellular interactions. Given the critical role of microglia in neurodegenerative diseases, integrating them into midbrain organoids offers a physiologically relevant platform for studying early neuroimmune contributions to Parkinson’s disease and testing potential therapeutic strategies (Cakir et al., 2022; Toh et al., 2023; Sabate-Soler et al., 2024).

In this study, we generated human midbrain assembloid models by integrating iPSC-derived microglia into midbrain organoids from both healthy individuals and PD patients with the *SNCA* triplication (3xSNCA). Upon integration into midbrain organoids, these microglia induced early and endogenous formation of phosphorylated α-synuclein (pS129) aggregates and PD-like pathology in the absence of exogenous fibril seeding. Notably, this pathology emerged by day 50 of culture, which is considerably earlier than previously observed in long-term models. Remarkably, this pathological process was observed not only in 3xSNCA assembloids but also in chimeric assembloids consisting of healthy midbrain organoids and PD specific microglia. This finding indicates that 3xSNCA microglia alone were sufficient to induce endogenous phospho-α-synuclein (pS129) pathology. These findings further highlight the crucial role of microglia in driving early-stage α-synuclein pathology and demonstrate the utility of midbrain assembloids as an advanced model for investigating the early neuroimmune contributions to Parkinson’s disease.

## Materials and methods

### Ethical approval

All work involving iPSCs was approved by the Ethics Review Panel (ERP) of the University of Luxembourg and the national Luxembourgish Research Ethics Committee (CNER, Comité National d’Ethique de Recherche) under CNER No. 201901/01 (ivPD) and No. 202406/03 (AdvanceOrg).

### iPSCs and NESCs

In this study, three human wildtype and one human *SNCA* triplication induced pluripotent stem cell (iPSC) lines were used, which are described in Supplementary Table 1. hiPSC were maintained in Essential E8 medium (Thermo Fisher Scientific, cat.no. A1517001) supplemented with 1% Penicillin-Streptomycin (Invitrogen, cat. no. 15140122) on Geltrex-coated plates (Corning, cat. no. 354277). They were passaged with Accutase (Sigma-Aldrich, cat. no. A6964) and cultured in E8 medium supplemented with 10 µM ROCK inhibitor Y-27632 (Merck Millipore, cat. no. 688000) for 24h after seeding (Gomez-Giro et al., 2019). Afterwards, the cells were cultured in E8 medium, with daily media exchanges.

Neuroepithelial stem cells (NESCs) were derived from iPSCs following to the protocol established by Reinhardt et al. (2013). Cells were cultured on Geltrex-coated plates in freshly prepared maintenance medium. The N2B27 base medium consisted of a 1:1 ratio of DMEM- F12 (Thermo Fisher Scientific, cat. no. 21331046) and Neurobasal (Thermo Fisher Scientific, cat. no. 10888022), supplemented with 1% GlutaMAX (Thermo Fisher Scientific, cat. no. 35050061), 1% penicillin/streptomycin (Thermo Fisher Scientific, cat. no. 15140122), 1% B27 supplement without Vitamin A (Life Technologies, cat. no. 12587001) and 2% N2 supplement (Thermo Fisher Scientific, cat. no. 17502001). For maintenance, the base medium was further supplemented with 150 µM ascorbic acid (Sigma-Aldrich, cat. no. A4544), 3 µM CHIR-99021 (Axon Medchem, cat. no. CT 99021), and 0.75 µM purmorphamine (Enzo Life Science, cat. no. ALX-420-045). NESc maintenance medium was replaced every other day.

### Microglia

Macrophage precursors were generated from iPSCs (van Wilgenburg et al., 2013) and subsequentially differentiated into microglia as previously described previously (Haenseler, Sansom, et al., 2017). For immunofluorescence staining, 150000 macrophage precursors were seeded onto a glass coverslip placed in 24-well plates (Thermo Fisher Scientific, cat. no. 142475). For flow cytometry and phagocytosis assays, 50000 macrophage precursors were plated in 96-well plates (Thermo Fisher Scientific, cat. no. 167008). Cells were cultured inmicroglia base medium composed of Advanced DMEM/F12 (Thermo Fisher Scientific, cat. no. 12634010), 1% Penicillin/Streptomycin (Invitrogen, cat. no. 15140122), 1% GlutaMAX™ (Thermo Fisher Scientific, cat. no. 35050061), 1% N2 (Thermo Fisher Scientific, cat. no. 17502001) and 50 μM 2-mercaptoethanol (Thermo Fisher Scientific, cat. no. 31350-010). The medium was further supplemented with 100 ng/ml GM-CSF (PeproTech, cat. no. 300-03) and 10 ng/ml IL-34 (PeproTech, cat. no. 200-34).

### Midbrain organoids

Midbrain organoids were generated from NESCs according to the protocol previously published by Monzel et al. (2017) and Zagare et al (2020) and cultured until day 50. NESCs were detached at 80% confluence with Accutase (Sigma-Aldrich, cat. no. A6964). Viable cells were counted with trypan blue, and to initiate sphero formation, 9000 live cells were seeded per well of a 96-well ultra-low attachment plate (faCellitate, cat. no. F202003) in 150 µl of NESC maintenance media. On day 2 of culture, the maintenance media was replaced with patterning media which consists of supplemented N2B27 base medium with 200 µM ascorbic acid (Sigma- Aldric, cat. no. A4544-100G), 500 µM dibutyryl-cAMP (STEMCELL Technologies, cat. no. 100- 0244), 10 ng/ml hBDNF (PeproTech, cat. no. 450-02-1mg), 10 ng/ml hGDNF (PeproTech, cat. no. 450-10-1mg), 1 ng/ml TGF-β3 (PeproTech, cat. no. 100-36E) and 1 µM purmorphamine (Enzo Life Science, cat. no. ALX-420-045). The next pattering media exchange was done on day 5 of organoid culture.

### Assembloids

Assembloids (ASBs) were cultured as described by Sabate-Soler et al (2022) with minor changes. On day 8 of culture midbrain organoid each midbrain organoid was coc-ultured with 50000 freshly harvested macrophage precursor cells. The culture medium was changed to the co-culture medium containing the microglia base medium supplemented with 100 ng/ml GM-CSF (PeproTech, cat. no. 300-03), 10 ng/ml IL-34 (PeproTech, cat. no. 200-34), 10 ng/ml BDNF (PeproTech, cat. no. 450-02) and 10 ng/ml GDNF (PeproTech, cat. no. 450-10). The medium was changed every 3-4 days, and the assembloids was kept for 50 days in culture. After the collection, the assembloids were either snap-frozen and stored at -80°C, or fixed with 4% formaldehyde (Millipore, cat. no. 1.00496.5000) for immunofluorescence staining.

iPSC, NESC and assembloids were regularly (once per month) tested for mycoplasma contamination using LookOut® Mycoplasma PCR Detection Kit (Sigma-Aldrich, cat. no. MP0035-1KT).

### Phagocytosis assay

For the phagocytosis assay, Zymosan A (S. cerevisiae) BioParticles™ (Thermo Fisher Scientific, cat. no. Z23373) or pHRodo™-Zymosan BioParticles™ Conjugate (Thermo Fisher Scientific, cat. no. P35364) were used. 10 µg/ml of particles were prepared in HBSS medium supplemented with 20 mM HEPES and 10% FCS. 50 µl of the microglial cell culture media was removed, and 50 µl of HBSS containing Zymosan A particles was added to the microglia. The cells were then incubated at 37°C for at least 30 minutes, washed twice with PBS, and analyzed via flow cytometry with the BD LSRFortessa (BD Biosciences, RRID:SCR_018655).

### Cytokine measurement

For intracellular cytokine measurement, microglia were stimulated for 24 hours with 100 ng/ml lipopolysaccharide (LPS) on day 14 of differentiation. During the final 6 hours of stimulation, BD GolgiPlug™ (BD Biosciences, cat. no. 555029) was added to the cell culture medium at a 1:1000 dilution. Microglia were then fixed and permeabilized using the BD Pharmingen™ Transcription Buffer Set (BD Biosciences, cat. no. 562574) following the manufacturer’s instructions. The microglia were washed twice in PBS, and immediately analysed in flow cytometer.

### Western Blot

For Western blot, 10 assembloids from three batches were pooled and lysed using RIPA buffer (Abcam, cat. no. ab156034) supplemented with cOmplete^TM^ Protease Inhibitor Cocktail (Sigma-Aldrich, cat. no. 116974980010) and Phosphatase Inhibitor Cocktail Set (Merck Millipore, cat. no. 524629). 100 µl of lysis buffer was added to the samples, disrupted by pipetting, and incubated on ice for 20 min. Lysates were sonicated for 10 cycles (30 sec on/30 sec off) using the Bioruptor Pico (Diagenode, RRID:SCR_023470). Samples were then centrifuged at 4°C for 30 min at 15,000 x *g*. The protein concentration was determined using the Pierce^TM^ BCA Protein Assay Kit (Thermo Fisher Scientific, cat. no. 56257423225) and samples were adjusted to the same concentration and boiled at 95°C for 5 min in denaturating loading buffer. Protein separation was achieved using SDS polyacrylamide gel electrophoresis (Bolt™ 4–12% Bis-Tris Plus Gel, Thermo Fisher Scientific, cat. no. 562574NW04120BOX) and transferred onto PVDF membrane using iBlot™ 2 Gel Transfer Device (Thermo Fisher Scientific, cat. no. IB21001). Membranes were fixed with 0.4% formaldehyde for 30 min at room temperature (RT), washed with PBS containing 0.02% Tween (PBST) and blocked for 1 h at RT in 5% BSA in PBS before incubating overnight (ON) at 4°C with the primary antibodies prepared in 5% BSA in PBS-T (Supplementary Table 2). The next day, membranes were washed three times for 5 min with PBST and incubated with DyLight^TM^ secondary antibodies at a dilution of 1:10,000 (anti-rabbit IgG (H+L) (DyLight 800 Conjugate, Cell Signaling, cat. no. 5151; RRID:AB_10697505) or anti-mouse IgG (H+L) (DyLight 680, Cell Signaling, cat. no. 5470, RRID:AB_10696895)) for 1 h at RT. Membranes were revealed in the Odyssey® Fc 2800 Imaging (LI-COR Biosciences, RRID:SCR_023227). Western blots were analysed using Image Studio Lite (LI-COR Biosciences, RRID:SCR_013715) software.

### Dot Blot for **α**-synuclein

150 µl assembloid media was collected from three batches, snap frozen and stored at -80°C. The supernatant was thawed on ice and centrifuged at 300 x *g* to sediment any cell debris remaining in the media. A Dot Blot Minifold I (Whatman, cat. no. 10447900) was used according to manufacturer guidelines. A nitro-cellulose membrane (Sigma-Aldrich, cat. no. GE10600001) was hydrated twice with 200 μl sterile PBS per well before sample loading. After sample run with the vacuum on, the membrane was washed with 200 μl sterile PBS per well. The membrane was retrieved and fixed for 10 min at 37°C, followed by a 1 min wash with PBS to hydrate the membrane. The membrane was incubated for 5 min with Revert 700 Total Protein Stain (LI-COR Biosciences, cat. no. 926-11016), followed by two quick washed with Milli-Q water, before revealing the membrane using the Odyssey® Fc 2800 Imaging System. The membrane was then destained for 5 min with the Revert destaining Solution (LI-COR Biosciences, cat. no. 926-11016) and washed once with PBS-T. Blocking and antibody incubation were done as previously described in the Western blotting section. Images were acquired with the Odyssey® Fc 2800 Imaging System. Images were analysed with Image Studio Lite and the relative α-synuclein amount was normalized to the total protein.

### Immunofluorescence staining of microglia

Macrophage precursors were plated on glass coverslips (150K/well) and cultured for 14 days. Then, mature microglia were fixed for 15 min with 4% formaldehyde (Sigma-Aldrich, cat. no. 100496) at RT and washed three times with PBS. The glass coverslips were permeabilized with 0.3% Triton X-100 in PBS for 15 min at RT. The cells were washed again three times with PBS, followed by blocking with 3% BSA (Carl Roth, cat. no. 80764) in PBS for 1h at RT. The microglia were then incubated with the primary antibodies (Supplementary Table 3) in 3% BSA, 0.3% Triton X-100 in PBS in a wet chamber ON at 4°C. Afterwards, the cells were washed three times with PBS before incubation with secondary antibody (Supplementary Table 3) in 3% BSA, 0.3% Triton X-100 in PBS in a wet chamber for 1h at RT. After three more PBS washed, the coverslips were mounted on a glass slide with Fluorormount-G® (Southern Biotech, cat. no. 0100-01).

### Immunofluorescence staining of assembloid sections

Assembloids were fixed ON at 4°C with 4% formaldehyde and then washed three times with PBS for 10 min at RT. For each condition, at least three to four assembloids were embedded in 3% low-melting point agarose (Biozym, cat. no. 840100). The solidified agarose blocks with the assembloid were sectioned using the vibrating blade microtome (Leica VT1000s, RRID:SCR_016495) at 60 μm. Sections were permeabilised with 0.5% Triton X-100 for 30 min at RT, followed by a short wash with 0.01% Triton X-100 in PBS. They were then blocked for 2h at RT with blocking buffer containing 2.5% BSA, 2.5% donkey serum, 0.01% Triton X-100 and 0.1% sodium azide in PBS. Primary antibodies (Supplementary Table 3) were diluted in blocking buffer and sections were incubated for 48-72h at 4°C on an orbital shaker. After incubation, the sections were washed with wash buffer (0.01% Triton X-100 in PBS) and incubated with secondary antibodies (Supplementary Table 3) in blocking buffer for 2h at RT and protected from light. The sections were then washed three times for 10 min at RT, rinsed with Milli-Q water and mounted as described by Nickels et al (2020).

### Proteostat® Aggresome Staining of assembloid sections

After fixation and sectioning, assembloid sections were permeabilized with 0.5% Triton X-100 and 3 mM EDTA (pH 8.0) in 1x Assay Buffer for 30 min at RT on an orbital shaker. After a wash with 0.01% Triton X-100 in PBS for 5 min at RT, they were blocked in blocking buffer as described in the immunofluorescence staining method. Primary antibodies (Supplementary Table 3) were diluted in blocking solution and incubated for 48-72h at 4°C on an orbital shaker, which was followed by one wash with 0.01% Triton-X 100 in PBS for 10 min at RT. The other two washes were done with PBS for 10 min at RT each. The secondary antibodies and the Proteostat® Aggresome Detection Reagent (Enzo, cat. no. ENZ-51035) were diluted (1:1000) in 1x Assay Buffer and incubated for 1h at RT on an orbital shaker. Sections were then washed three times with PBS for 10 min at RT each. They were once washed with Milli-Q water before mounting.

### Thioflavin S staining of assembloid sections

Thioflavin S staining was performed based on the method described by Stojkovska and Mazzulli (2021). Assembloid sections were fixed and sectioned as previously described and permeabilized with 0.3% Triton X-100 in PBS for 30 min at 4°C on an orbital shaker. They were then blocked for 30 min at RT on an orbital shaker with 2.5% BSA, 2.5% normal goat serum and 0.3% Triton X-100 in PBS. Assembloid sections were then incubated with the primary antibodies (Supplementary Table 3) in blocking solution and incubated ON at 4°C on an orbital shaker. Afterwards three washes with 0.01% Triton X-100 in PBS for 10 min each at RT were performed followed by incubation of secondary antibody in 0.01% Triton X-100 in PBS (Supplementary Table 3) for 2h at RT and protected from light. It is important to note that the Hoechst 33342 dye is not added in this staining protocol. Sections were then washed again three times for 10 min at RT with the washing buffer before incubation with the fresh Thioflavin S solution which consists of 0.05% Thio S (Sigma-Aldrich, cat. no. T1892) w/v in 50% EtOH/Milli-Q water. Afterwards the assembloid sections were washed two times with 50% EtOH for 20 min at RT, before a consecutive wash with 80% EtOH for 20 min at RT. Finally, a last wash was performed with the washing buffer before rinsing the sections with Milli-Q water and mounting.

### Image acquisition

For high-content imaging, mounted assembloids were imaged with the Yokogawa CellVoyager CV8000 microscope (Yokogawa, RRID:SCR_023270). A 4x pre-scan identified the wells containing assembloid sections, which enabled the creation of masks to outline the assembloids. These masks guided the selection of the field for imaging at different wavelengths with a 20x objective.

Qualitative images were acquired with the Zeiss LSM 710 confocal laser scanning microscope (Zeiss, RRID:SCR_018063) equipped with 20x or 63x objectives. Three-dimensional reconstructions of confocal Z-stacks were created using Imaris software (Bitplane, RRID:SCR_007370). Representative images were cropped, rotated, or rescaled for visualisation using Adobe Photoshop (version 26.6.1, Adobe, RRID:SCR_014199) and Illustrator (version 29.5.1, Adobe, RRID_SCR_010279). All original, unedited images are available at https://doi.org/10.17881/syhp-k282.

### Image processing, and analysis

For quantitative image analysis, at least two sections from three assembloids, each derived from three independent batch generations, were analysed in MATLAB (2021a, Mathworks, RRID:SCR_001622). A custom image-analysis algorithm, as previously described by Bolognin et al. and Monzel et al. (2018, 2020), was used. This algorithm merges overlapping sections into mosaic images, performs smoothing, combines colour channels and filters out small objects.

Marker-specific masks were created based on pixel intensity thresholds and refined to quantify marker intensities in 3D space (voxels).

### RNA extraction, library preparation and sequencing

Total RNA of microglia was extracted from microglia using RNeasy Mini Kit (Qiagen, cat. no. 74104) according to the manufacturer’s protocol. For each cell line, RNA was isolated from three pooled wells of a 6-well plate, with each well containing 1x10^6^ macrophage precursors. Messenger RNA (mRNA) was purified from total RNA with the poly-T-oligo-attached magnetic beads. Following RNA fragmentation, first-strand cDNA was synthesized using random hexamer primers. Second-strand cDNA synthesis was performed using either dUTP (for directional library) or dTTP (for non-directional library). Libraries were sequenced on an Illumina platform by Novogene’s sequencing service.

### Transcriptomic analysis

RNA sequencing data from microglia were pre-processed with the Galaxy server (version 23.2.rc1, RRID:SCR_006281), following established Galaxy training tutorials (Hiltemann et al., 2023, Doyle et al., 2024). Sequencing reads were aligned to the human reference genome (hg38) using HISAT2 (RRID:SCR_015530), and read counts were generated from the resulting BAM files using the *featureCounts* function. Differential gene expression analysis was conducted in R (version 4.4.2., RRID:SCR_001905) using the "DESeq2" software package (version 1.42.1, RRID:SCR_000154) (Love et al., 2014). P-values were corrected for multiple hypothesis testing according to the Benjamini-Hochberg method (Benjamini & Hochberg, 2005). Pathway enrichment analysis was performed using the EnrichR tool (RRID:SCR_001575) (Kuleshov et al., 2016), based on the expressed genes identified by DESeq2. Two gene set libraries were used for enrichment analysis: “Gene Ontology (GO) Biological Process 2023” (Ashburner et al., 2000, Aleksander et al., 2023) and “Kyoto Encyclopedia of Genes and Genomes (KEGG) Human 2021” (Kaneshisa et al., 2000, 2019, 2025).

### Statistical analysis

The data in this manuscript was processed, analysed and visualised in both GraphPad Prism (GraphPad version 10.2.3, RRID:SCR_002798) or the R software (R version 4.4.2, RRID:SCR_001905). Statistical significance of non-normally distributed data was tested with a non-parametric Mann-Whitney *U* rank sum test for comparison between the two conditions (WT microglia and 3xSNCA microglia; WT ASB and 3xSNCA ASB; WT ASB and WT-3xSNCA ASB).

Each data point in the graphs corresponds to the data from one cell line for one individual differentiation experiment. Significance asterisks in the figure legends represent *p < 0.05, **p < 0.01, ***p < 0.001, ****p < 0.0001 . When data was found not significant, it is not specifically stated in the figures and is expressed as ns, not significant. Error bars represent mean + standard deviation (SD). Detailed information on the number (n) of samples, replicates and batches is added to the individual Figure legends.

## Results

### 3xSNCA microglia express α-synuclein, exhibit reduced phagocytosis and show altered inflammatory activity

In this study, we used three different iPSC lines from healthy individuals and the 3xSNCA cell line derived from a PD patient with a *SNCA* triplication (Supplementary Table 1). A *SNCA* knockout cell line (KO) was also included for validation of α-synuclein antibody specificity. To investigate the mature microglia, we derived macrophage precursors from human iPSCs that were differentiated into microglia for 14 days as previously described by Haenseler, Sansom, et al. (2017) (Figure 1a). Immunofluorescence staining confirmed expression of macrophage-specific markers, such as CD45, PU.1, and IBA1, and microglia markers, including TMEM119 and P2RY12 (Figure 1b, Supplementary Figure 1a-c, Supplementary Figure 2a). After confirming the microglial cell identity, we stained the mature microglia with an anti-α-synuclein antibody and showed that the protein is expressed in both WT and 3xSNCA microglia, but not visible in the KO microglia (Figure 1b, Supplementary Figure 2a). We could also determine a nuclear localization of the α-synuclein protein in both the WT and 3xSNCA microglia (Figure 1b, Supplementary Figure 2a). Additionally, extracellular α-synuclein release was measured by dot blot and shows a significant increase in the 3xSNCA microglial medium (Figure 1c).

**Figure 1:**
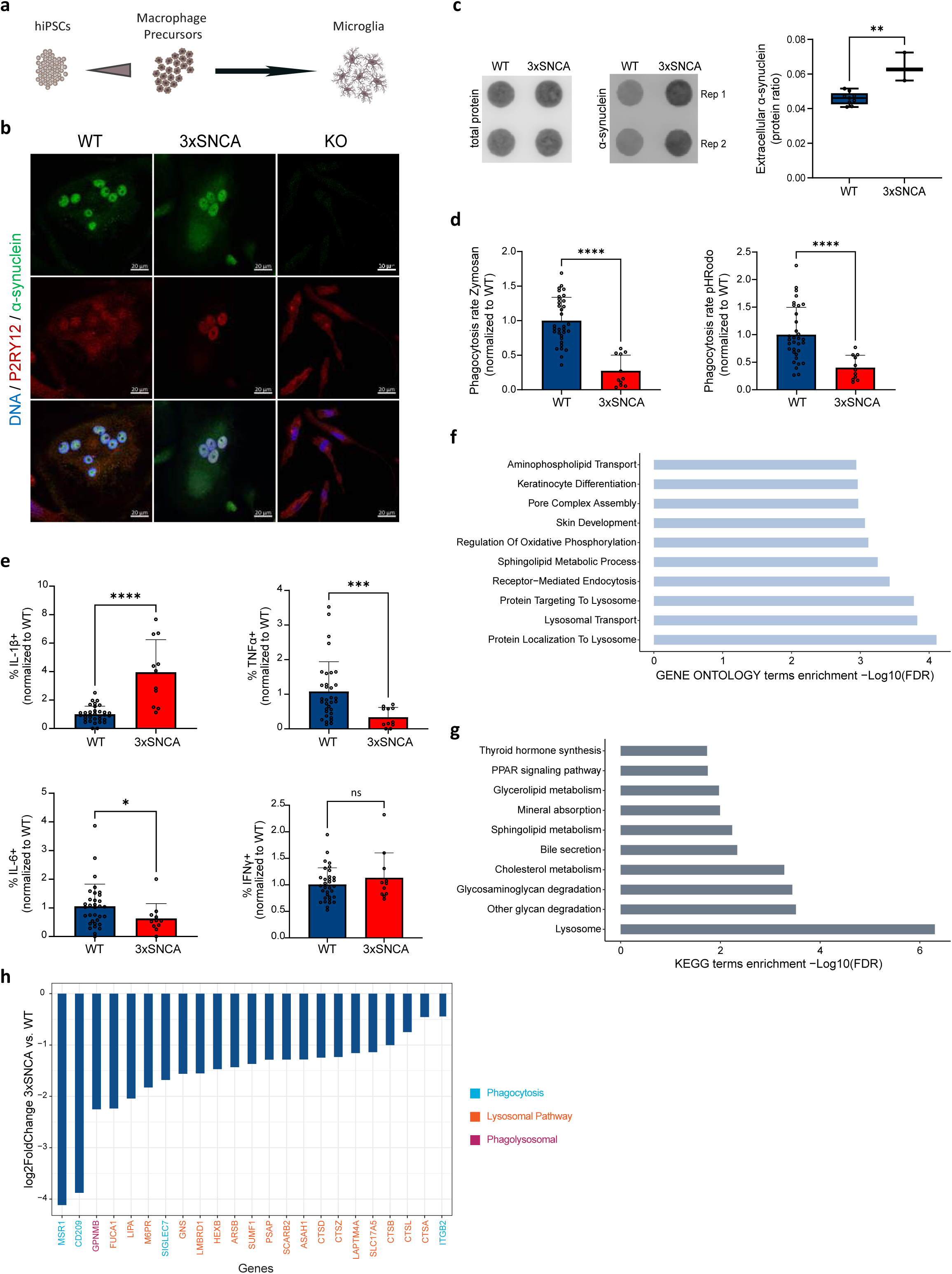
3xSNCA iPSC-derived microglia express α-synuclein and functional deficits. (a) Schematic representation of iPSC-derived microglia via macrophage precursors (pMacPre). (b) Representative immunofluorescence staining of mature (day 14) WT, 3xSNCA and KO microglia for the microglial marker P2RY12 (Purinergic Receptor P2Y12), total α-synuclein and a DNA marker (Hoechst 33342) (scale bar 20 μm, 63x). (c) Representative dot blot (right panel) of total protein (Revert700) and anti-α-synuclein antibody showing higher levels of extracellular α-synuclein released by 3xSNCA microglia compared to WT microglia (left panel) at day 14. Data is represented as boxplots and shows the mean of two technical replicates normalized to the total protein (Revert700) of three independent microglia batches (n=3). Mann-Whitney *U* test; **p < 0. 01. Dot blots are cropped from original images found in Supplementary original blots. (d) Flow cytometry analysis of microglial phagocytosis in WT and 3xSNCA microglia using fluorescently labelled Zymosan beads (left panel) and pH sensitive fluorescent Zymosan beads (pHRodo) (right panel). Data is represented as bar plots and shows three independent batches (n=3) normalized to the mean of the controls per batch. Mann-Whitney *U* test; ****p < 0.0001. (e) Flow cytometry analysis of intracellular IL-1β (upper left panel), TNF-α (upper right panel), IL-6 (lower left panel) and IFN-γ (lower right panel) in WT and 3xSNCA microglia after 24h stimulation with LPS. Data is represented as bar plots and shows three independent batches (n=3) normalized to the mean of the controls per batch. Mann-Whitney *U* test; *p<0.05, ***p<0.001, ****p < 0.0001. (f) GO pathway enrichment analysis of the DEGs (p.adjusted <0.01) between 3xSNCA and WT samples, showing the top 10 significantly enriched pathways based on EnrichR results. (g) KEGG pathway enrichment analysis of the DEGs (p.adjusted <0.01) between 3xSNCA and WT samples, showing the top 10 significantly enriched pathways based on EnrichR results. (h) Log_2_ fold changes (FC) of predefined genes associated with phagocytosis (blue), the lysosomal pathway (orange), or both pathways (phagolysosomal, magenta).

Next, we assessed the phagocytic capacity of the mature microglia. Incubation with Zymosan particles showed a significant decrease in Zymosan particle uptake (Figure 1d, left panel) along with a reduced pHRodo fluorescence rate (Figure 1d, right panel) in the 3xSNCA microglia, indicating an impairment in both the internalization and phagosomal acidification process, suggesting a deficiency in effective phagocytosis. Haenseler, Zambon et al. (2017) similarly reported accumulation of α-synuclein along with impaired phagocytic function in iPSC-derived 3xSNCA macrophages, supporting our findings.

Chronic activation of microglia leads to excessive release of cytokines such as IL-1, TNF-α, IL- 6, and IFN-γ, creating a toxic environment for neurons (Tansey et al., 2022). Previous studies have shown that elevated α-synuclein levels induce a pro-inflammatory response in microglia (Zhang et al., 2005; Klegeris et al., 2008; Ferreira & Romero-Ramos, 2018). Notably, we wanted to determine the impact of the SNCA triplication on microglial cytokine production in our WT and 3xSNCA microglia. Flow cytometry analysis showed that IL-1β, a key mediator of the inflammatory response, was significantly upregulated in the 3xSNCA microglia compared to the healthy controls (Figure 1e, upper left panel). Interestingly, both TNF-α and IL-6 were significantly downregulated, and no change was seen in IFN-γ (Figure 1e). Together these results indicate that 3xSNCA microglia exhibit increased α-synuclein expression, impaired phagocytosis, and altered inflammatory activity, highlighting their potential role in neuroinflammation and Parkinson’s disease pathology.

### Transcriptomic data analysis reveals lysosomal dysfunction in 3xSNCA microglia

To further elucidate the molecular mechanisms underlying these phenotypic alterations, we employed RNA sequencing to comprehensively analyse transcriptomic differences between WT and 3xSNCA microglia. By analysing transcriptomic data, we aimed to identify key dysregulated pathways and genes that may contribute to the observed cellular dysfunctions, providing further insights into the role of α-synuclein in microglial pathology. We identified 149 differentially expressed genes (DEGs) in 3xSNCA microglia compared to the three WT cell lines, using a threshold of adjusted p-value (FDR) <0.01 (Supplementary Figure 2b). Unsupervised hierarchical clustering based on normalised gene counts of top significant DEGs involved in the inflammasome pathway further highlighted clear separation between the two groups and confirms our findings on altered inflammatory activity (Supplementary Figure 2c). Gene Ontology pathway enrichment analysis of the p-value <0.01 significant DEGs between 3xSNCA and WT samples revealed that the lysosomal pathway is dysregulated, including protein localization to lysosome, lysosomal transport, and protein-targeting to lysosome, as the top enriched pathways (Figure 1f). Additionally, KEGG (Kyoto Encyclopedia of Genes and Genomes) enrichment analysis supported these results and identified “Lysosome” as the most enriched pathway, followed by metabolic pathways such as the lipid metabolism and carbohydrate metabolism (Figure 1g). The enrichment analysis suggests that lysosomal dysfunction and metabolic dysregulation are key processes affected at the transcriptomic level. Looking at the expression of individual genes of the phagocytic and lysosomal pathway, we could confirm that the key genes of these pathways are downregulated in the 3xSNCA microglia (Figure 1h). Our findings highlight disruptions in cellular clearance mechanisms and metabolic balance, emphasizing the critical role of lysosomal and phagocytic pathways in neurodegeneration.

### 3xSNCA microglia integrated into midbrain organoids display morphological differences

After describing the phenotypic and transcriptomic changes in 3xSNCA microglia, we sought to investigate these cells in a physiologically relevant context. To achieve this, we integrated the microglia into midbrain organoids following the protocol described by Sabate-Soler et al. (2022) (Figure 2a). This approach allowed us to study microglial behaviour and interactions within a three-dimensional cellular environment that closely mimics the complexity of the human brain. Previous studies have shown that changes in microglial function, such as increased reactivity, enhanced pro-inflammatory response, impaired phagocytosis, altered gene expression, and exacerbation of α-synuclein pathology, contribute to the chronic neuroinflammation and neurodegeneration observed in PD (Thi et al., 2024; Zhu et al., 2022).

**Figure 2:**
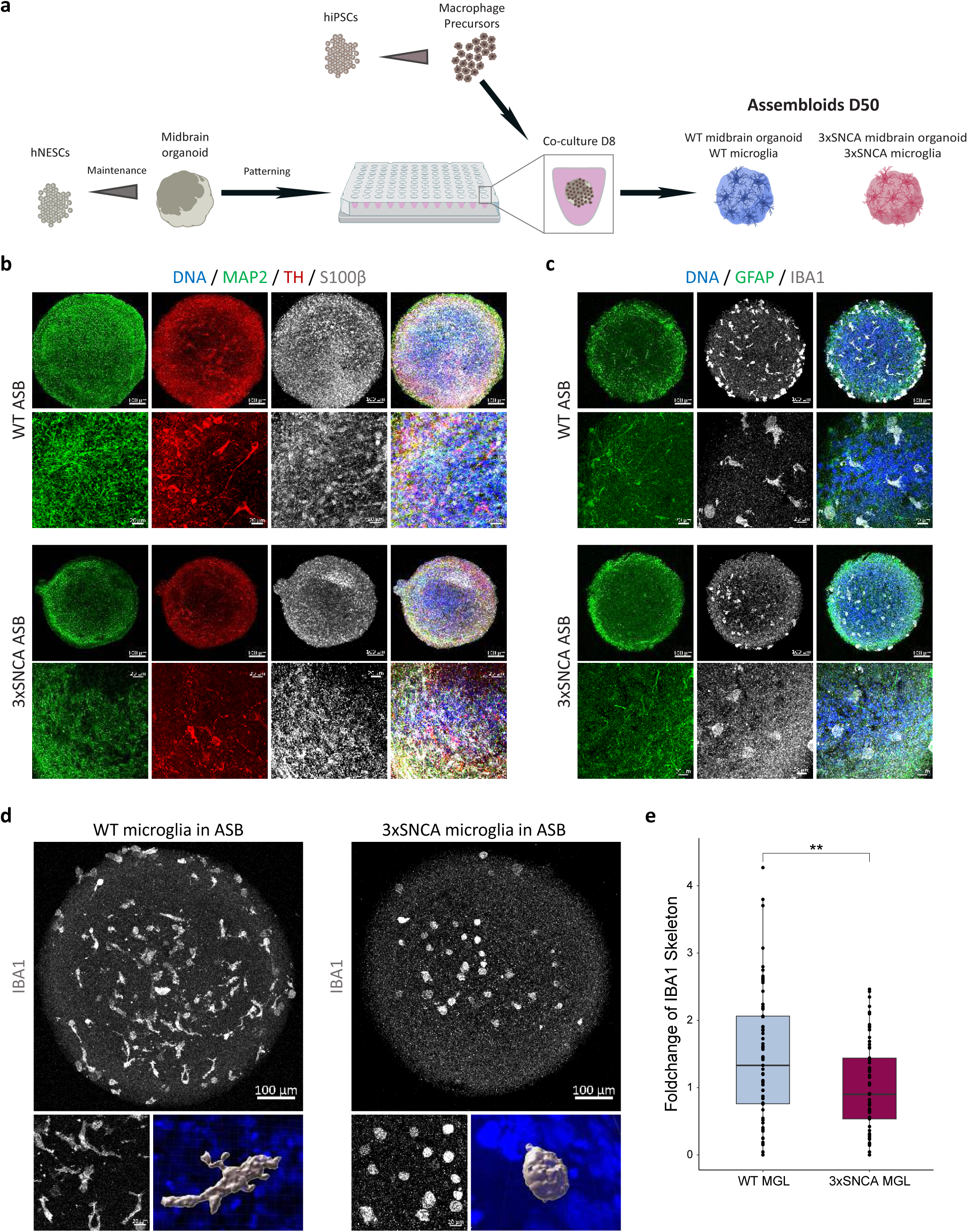
3xSNCA microglia integrated into midbrain organoids display morphological differences. (a) Schematic representation of microglia integration into the midbrain organoid. At day 8 of culture, macrophage precursor cells are added to the midbrain organoid and the assembloids are culture until 50 days of culture. WT microglia are integrated into the WT midbrain organoid, and 3xSNCA patient-derived microglia into the 3xSNCA midbrain organoid. (b) Representative immunofluorescence staining of WT assembloids (WT ASB) and 3xSNCA assembloids (3xSNCA ASB) with the dopaminergic neuron marker TH (tyrosine hydroxylase), the neuronal marker MAP2 (Microtubule-associated protein 2), the astrocyte marker S100β (S100 calcium-binding protein B) and Hoechst 33342. (c) Representative immunofluorescence staining of WT ASBs and 3xSNCA ASBs with the microglial marker IBA1 (Ionized calcium- binding adaptor molecule 1), the astrocyte marker GFAP (Glial fibrillary acidic protein) and Hoechst 33342 (scale bar 100 μm, 20x; 20 μm, 63x). (d) Representative immunofluorescence staining of WT and 3xSNCA microglia integrated into midbrain organoids stained with the microglial marker IBA1 and Hoechst 33342 (scale bar 100 μm, 20x; 20 μm, 63x). 3D reconstruction with IMARIS software showing the microglial morphology of the WT and 3xSNCA microglia in the midbrain organoid. (e) High-content automated image analysis of immunofluorescence stainings showing the foldchange of IBA1 skeleton in microglia. Data is represented as boxplots and show three to four independent batches (n=3-4) normalized to the mean IBA1 skeleton value within each batch. Mann-Whitney *U* test; **p < 0.01.

First, we characterized 50-day-old assembloids (ASBs) to ensure that they expressed the relevant cell types of the midbrain, showing the presence of mature neurons (MAP2 – microtubule associated protein 2), dopaminergic neurons (TH – Tyrosine Hydroxylase) and astrocytes (S100β – S100 calcium binding protein B) (Figure 2b). Staining with the microglial marker IBA1 (Ionized calcium binding adaptor molecule 1) and the astrocyte marker GFAP (glial fibrillary acidic protein) completed the characterization of the assembloids (Figure 2c). Interestingly, when examining the morphology of the integrated microglia, we observed striking differences between WT and 3xSNCA conditions. WT microglia in WT midbrain organoids displayed a more typical ramified morphology, characteristic of resting or surveillant microglia in a healthy state (Figure 2d, left panel, and 2e). In contrast, 3xSNCA microglia in 3xSNCA midbrain organoids exhibited an ameboid morphology, indicative of an activated state (Figure 2d, right panel, and 2e). These morphological distinctions, suggest that the 3xSNCA mutation leads to a more activated microglial phenotype, within the complex 3D organoid environment.

### 3xSNCA assembloids exhibit elevated α-synuclein and phospho-α-synuclein (pS129) levels

Given that the SNCA triplication leads to increased expression of α-synuclein and is associated with early-onset Parkinson’s disease (Singleton et al., 2003), we investigated both total α- synuclein and its phosphorylated form (pS129) to assess disease-relevant phenotypes in 3xSNCA assembloids. α-synuclein protein levels were evaluated by Western Blot, showing an expected significant increase in the 3xSNCA assembloids (Figure 3a and 3b, left panel). The specificity of the α-synuclein antibody has been previously confirmed by the lack of signal in SNCA KO midbrain organoids (Muwanigwa et al., 2024). Consistent with its role as a hallmark of pathology, pS129 was also significantly elevated in 3xSNCA assembloids (Figure 3a and 3b, right panel), in agreement with previous findings. α-synuclein and pS129 staining of assembloids validated these findings by showing significantly increased intensities in the 3xSNCA assembloids (Figure 3c and 3d). Additionally, dot blot analysis revealed enhanced extracellular α-synuclein release (Figure 3e).

**Figure 3:**
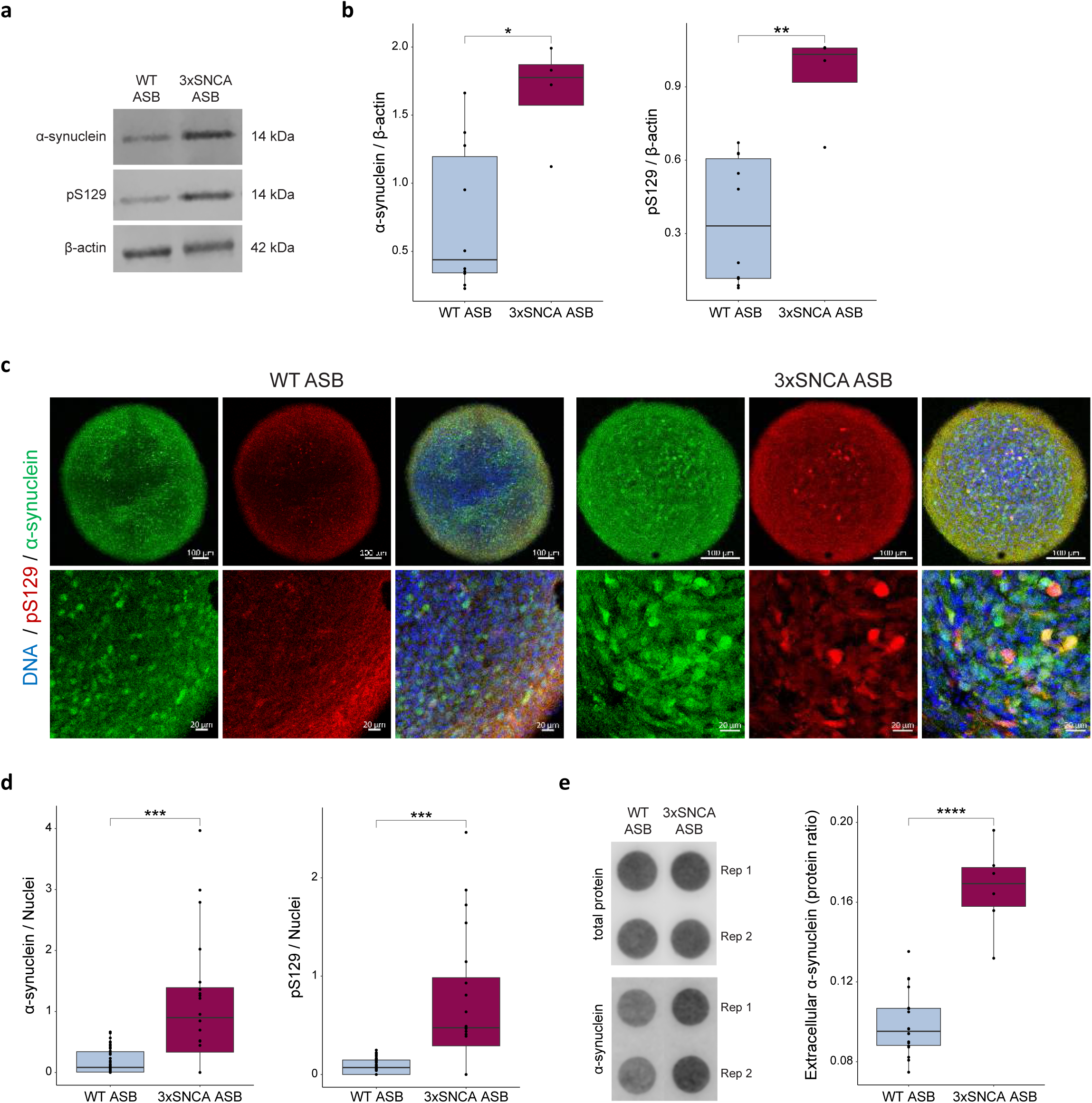
3xSNCA assembloids exhibit elevated total α-synuclein and phospho-α-synuclein (pS129) levels. (a) Representative immunoblots of α-synuclein, phospho-α- synuclein (pS129) and β-actin of day 50 WT and 3xSNCA ASBs. Western blots are cropped from original images found in Supplementary original blots. (b) Quantification of α-synuclein and phospho-α-synuclein (pS129) protein levels. Data is shown as boxplots and represents a summary of three to four independent batches (n=3-4) per cell line. Values are normalized to β- actin levels. Mann-Whitney *U* test; *p < 0.05, **p < 0.01. (c) Representative immunofluorescence staining of WT and 3xSNCA ASBs with total α-synuclein, pS129 and Hoechst 33342 (scale bar 100 μm, 20x; 20 μm, 63x). (d) High-content automated image analysis of immunofluorescence stainings showing the foldchange of α-synuclein (left panel) and pS129 (right panel) normalized by total nuclei. Data is shown as boxplots and represents three to four independent batches (n=3-4) normalized to the mean of the controls per batch. Mann-Whitney *U* test; ***p < 0.001. (e) Representative dot blot (left panel) of total protein (Revert700) and anti-a-synuclein antibody showing higher levels of extracellular α-synuclein released by 3xSNCA ASBs (right panel) at 50 days. The Boxplots represents three to four independent batches (n=3-4) normalized to the total protein (Revert700). Mann-Whitney *U* test; ****p < 0.0001. Rep = replicate. Dot blots are cropped from original image found in Supplementary original blots.

### 3xSNCA microglia promote endogenous formation of phospho-α-synuclein (pS129) pathology in assembloids

To evaluate whether the PD patient-specific model recapitulates key pathological features, we examined α-synuclein aggregation. We particularly focused on phosphorylation at serine 129 (pS129). This post-translational modification is a well-established marker of Lewy-body pathology and neuronal dysfunction (Spillantini et al., 1997; Gallegos et al., 2015; Samuel et al., 2016; Calabresi, Mechelli, et al., 2023). Notably, pS129 accounts for around 90% of α-synuclein within Lewy bodies, compared to only around 4% in healthy brains (Xu et al., 2015), and is associated with fibril formation, toxicity, and disease progression (Henderson et al., 2019).

Previous studies demonstrated that pS129 antibodies can detect diverse α-synuclein aggregate structures *in vitro*, specifically in neurons treated with preformed fibrils (PFFs) (Lashuel et al., 2022). Building on this, we assessed pS129-positive aggregates in 3xSNCA assembloids and were able to recapitulate these findings in the *in vitro* assembloid model without the addition of exogenous PFFs. Consistent with Lashuel’s observations, we identified distinct pS129-positive α-synuclein structures in our 3xSNCA assembloids, including Ring-like (Figure 4a, upper panel), Filamentous-like (middle panel), and Dense-like (lower panel) formations. To confirm the specificity of the observed pS129 pathology, we tested several anti-pS129 antibodies, including D1R1R (Figure 4a), MJF-R13 (8-8) (Supplementary Figure 3a), MJFR-14-6-4-2 (Supplementary Figures 3b and 3c). All consistently labelled pathological α-synuclein structures, supporting the reliability of our findings. Together, these results demonstrate that the 3xSNCA assembloid model spontaneously develops diverse α-synuclein pathological structures, further validating its relevance for studying synucleinopathies.

**Figure 4:**
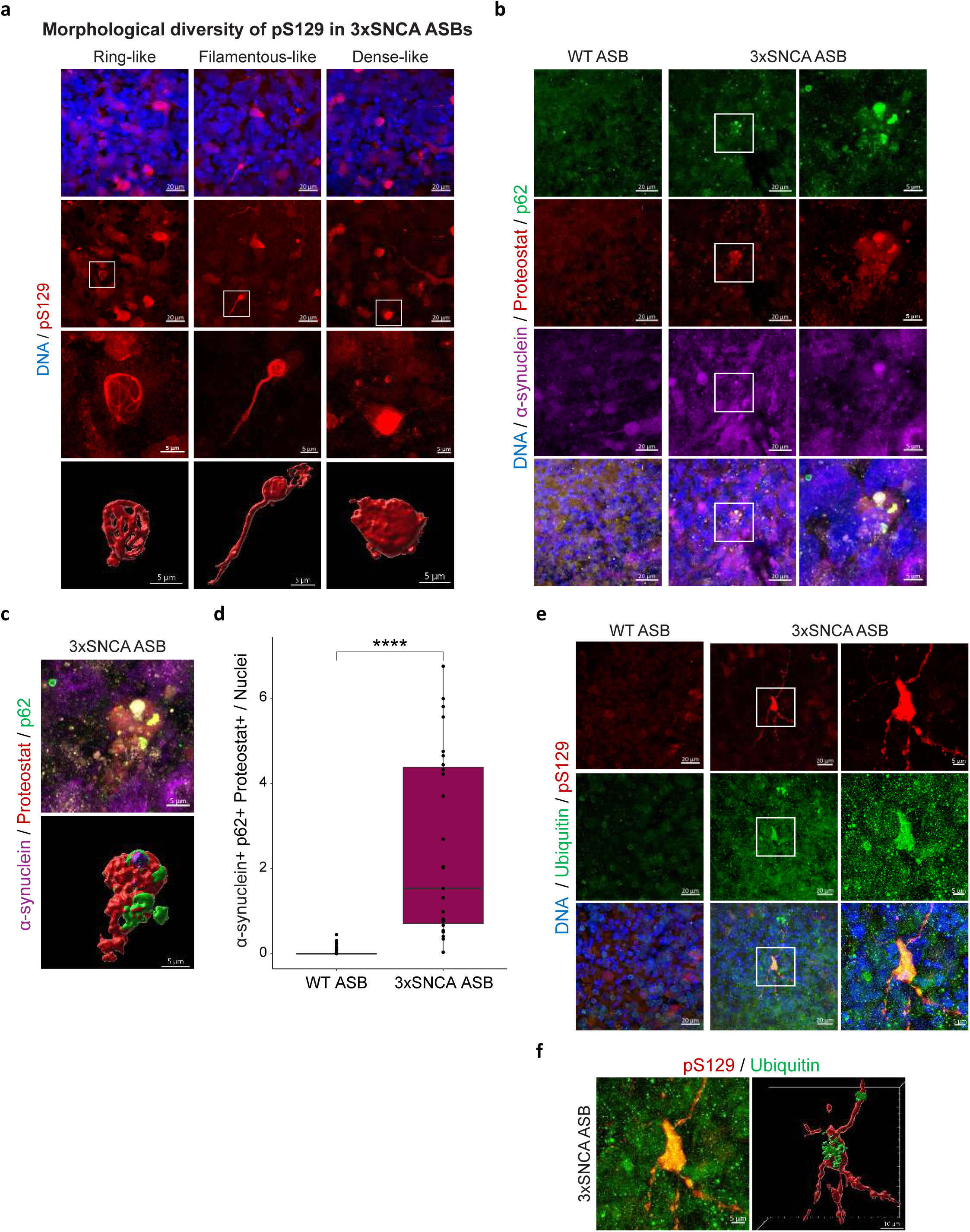
3xSNCA microglia promote endogenous formation of phospho-α-synuclein (pS129) pathology in assembloids. (a) Representative image of the morphological diversity of pS129 in 3xSNCA ASBs. The ring-like (upper panel), the filamentous-like (middle panel) and the dense-like (lower panel) can be distinguished with the pS129 marker (scale bar 20 μm, 63x). Zoom in of inset shows the pS129 structures (scale bar 5 μm). 3D reconstruction with IMARIS software shows the pS129 structural morphology. (b) Representative immunofluorescence staining of WT and 3xSNCA ASBs with total α-synuclein, Proteostat, p62 (SQSTM1) and Hoechst 33342 (scale bar 20 μm, 63x; 5 μm, zoom in of inset). (c) *Top panel:* The same representative image of 3xSNCA ASB as shown in (b), displayed here with total α-synuclein, Proteostat, and p62 (SQSTM1) markers (Hoechst omitted). *Lower panel:* 3D reconstruction of this image using IMARIS software (scale bar, 5 μm; zoom-in of inset). (d) High-content automated image analysis of immunofluorescence stainings showing the foldchange of α- synuclein+, p62+, Proteostat+ normalized by total nuclei. Data is shown as boxplots and represents three independent batches (n=3) normalized to the mean of the controls per batch. Mann-Whitney *U* test; ****p < 0.0001. (e) Representative immunofluorescence staining of WT and 3xSNCA ASBs with pS129, Ubiquitin and Hoechst 33342 (scale bar 20 μm, 63x; 5 μm, zoom in of inset). (f) *Left panel:* The same representative image of 3xSNCA ASB as shown in (e) displayed here with pS129 and Ubiquitin markers (Hoechst omitted). *Right panel:* 3D reconstruction of this image using IMARIS software (right panel) of 3xSNCA with (scale bar 5 μm, zoom in of inset).

Moreover, we used a comprehensive panel of markers to further validate the presence of pathological forms of α-synuclein in 3xSNCA assembloids originally detected with anti-pS129 antibodies (Canerina-Amaro et al., 2019). We observed a significant increase in the co- localization of α-synuclein with p62 (SQSTM1) and Proteostat in 3xSNCA assembloids (Figure 4b-d), indicating a substantial burden of α-synuclein–specific pathological aggregates. p62 is an autophagy adaptor protein frequently found in Lewy bodies, while Proteostat binds β-sheet–rich misfolded proteins, serving as a general marker of proteostasis disruption (Kuusisto et al., 2003; Rocha et al., 2018). Although p62 and Proteostat signals are also present in WT assembloids (Supplementary Figure 3d and 3e), the extent of their co-localisation with α-synuclein is substantially higher in the 3xSNCA assembloids (Figure 4d). To further support this finding, we confirmed additional markers through co-localization of pS129 with ubiquitin, a key component of the protein degradation machinery that accumulates at sites of misfolded protein aggregation (Figures 4e and 4f). These results suggest that elevated α-synuclein expression in the mutant assembloids drives the formation of misfolded protein aggregates.

Additionally, staining with Thioflavin S, which specifically labels β-sheet-rich amyloid structures, revealed co-localization with both α-synuclein and pS129 (Supplementary Figure 3b and 3c), indicating that these aggregates acquire amyloid-like conformations characteristic of mature pathogenic inclusions (Lee et al., 2002). In addition, co-localization of α-synuclein with the mitochondrial outer membrane marker TOM20 (Supplementary Figure 3f and 3g) suggests a potential interaction of aggregates with mitochondria, consistent with previous findings that link α-synuclein pathology to mitochondrial dysfunction (Di Maio et al., 2016). This enhanced co- localization provides strong evidence for the formation of α-synuclein aggregates in the 3xSNCA assembloid model, supporting the presence of pathological α-synuclein species and potentially indicating impaired protein degradation pathways.

### 3xSNCA microglia are sufficient to induce pS129 pathology in WT midbrain organoids

To investigate whether 3xSNCA microglia are sufficient to induce α-synuclein pathology in an otherwise healthy environment, we generated chimeric assembloids by integrating 3xSNCA mutant microglia into WT midbrain organoids (WT-3xSNCA ASB) (Figure 5a). Immunofluorescence analysis revealed a significant increase in both total α-synuclein and pS129 signal intensity in WT-3xSNCA assembloids compared to WT assembloids (Figure 5b), indicating that PD microglia alone are sufficient to induce pathology. Nevertheless, while the induction of pathology is clear, it is lower than what we saw before in organoids carrying the 3xSNCA mutation themselves (Figure 3d). We further visualized disease-relevant α-synuclein pathology in WT–3xSNCA assembloids, including compact and filamentous-like formations (Figure 5c), similar to those described in Lashuel et al (2020). Notably, these structures were detectable upon integration of 3xSNCA microglia into all three WT midbrain organoid lines. Aggregation was validated through co-localization of α-synuclein with p62, whose accumulation indicates impaired autophagic clearance, and Proteostat, a marker of misfolded protein aggregates (Figure 5d–e, Supplementary Figure 4a and 4b). To strengthen this finding, we confirmed additional markers by co-localization of pS129 with ubiquitin (Supplementary Figures 4c and 4d), and α-synuclein with the mitochondrial marker TOM20 (Supplementary Figures 4e and 4f), supporting the presence of Lewy body–like structures and mitochondrial involvement. Together, these results demonstrate that 3xSNCA microglia are sufficient to initiate α-synuclein pathology in an otherwise WT environment and highlight their emerging role as drivers of Parkinson’s disease pathogenesis.

**Figure 5:**
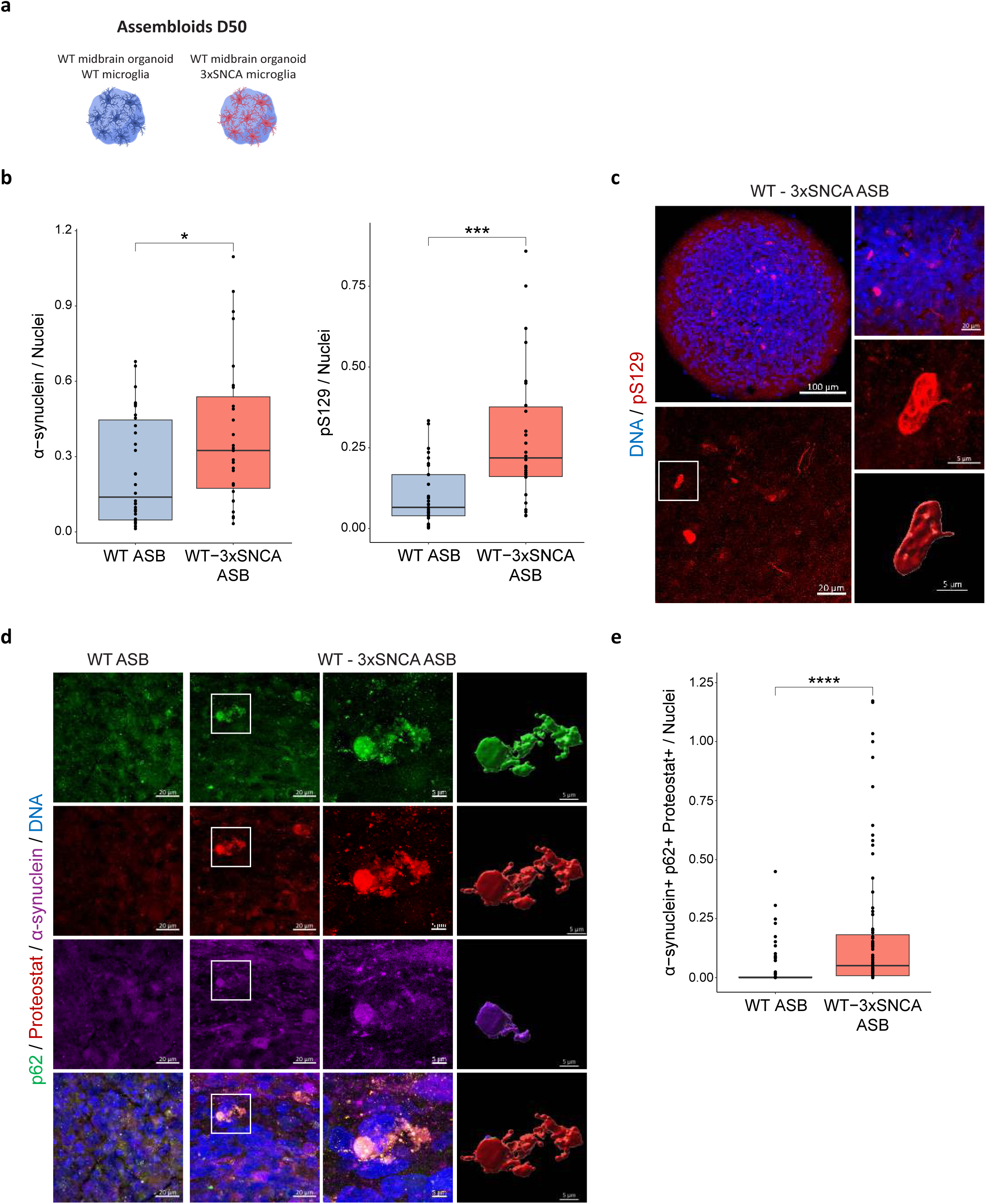
3xSNCA microglia are sufficient to form pS129 pathology in WT midbrain organoids. (a) Schematic representation of the chimeric assembloid model. WT microglia are integrated into the WT midbrain organoid (WT ASB), and 3xSNCA microglia into the WT midbrain organoid (WT-3xSNCA ASB). (b) High-content automated image analysis of immunofluorescence stainings showing the foldchange of α-synuclein (left panel) and pS129 (right panel) normalized by total nuclei. Data is shown as boxplots and represents three to four independent batches (n=3-4) normalized to the mean of the controls per batch. Mann-Whitney *U* test; *p < 0.05, ***p < 0.001. (c) Representative image of pS129 structures in 3xSNCA-WT ASBs. Sections were stained with pS129 marker and Hoechst 33342 (scale bar 100 μm, 20x; 20 μm, 63x; 5 μm, zoom in of inset). 3D reconstruction with IMARIS software shows the pS129 structural morphology. (d) Representative immunofluorescence staining of WT and WT-3xSNCA ASBs with total α-synuclein, Proteostat, p62 (SQSTM1) and Hoechst 33342 (scale bar 20 μm, 63x; 5 μm, zoom in of inset) and 3D reconstruction with IMARIS software (scale bar 5 μm, zoom in of inset). (e) High-content automated image analysis of immunofluorescence stainings showing the foldchange of α-synuclein+, p62+, Proteostat+ normalized by total nuclei. Data is shown as boxplots and represents three independent batches (n=3) normalized to the mean of the controls per batch. Mann-Whitney *U* test; ****p < 0.0001.

## Discussion

The endogenous formation of phosphorylated α-synuclein (pS129) remains a major challenge in PD modelling. Most *in vitro* systems and *in vivo* mouse models require the addition of preformed α-synuclein fibrils (PFFs) to induce Lewy body–like pathology (Luk et al., 2009; Volpicelli-Daley et al., 2011; Osterberg et al., 2015; Mahul-Mellier et al., 2020; Awa et al., 2022). In this study, we present a robust and reproducible patient-specific human midbrain assembloid model that recapitulates hallmark features of α-synuclein pathology, including the spontaneous formation of pS129-positive aggregates in the absence of exogenous fibrils. Our system integrates 3xSNCA iPSC-derived microglia into midbrain organoids, providing a physiologically relevant, tissue-like environment for studying early PD pathology.

Already, the 3xSNCA microglia alone exhibited impaired phagocytic function, altered inflammatory signalling, and revealed significant alterations in lysosomal pathway and metabolic processes consistent with previous work on iPSC-derived macrophage precursors from SNCA triplication patients (Haenseler, Zambon et al., 2017). Upon integration into midbrain organoids, these microglia showed an ameboid, activated morphology, consistent with their pro-inflammatory profile and loss of homeostatic surveillance. These phenotypes were retained in 3D assembloids and are likely contributors to α-synuclein accumulation and impaired clearance (Choi et al., 2020; Bido et al., 2021; Eo et al., 2024).

Previous studies have consistently reported increased α-synuclein and progressive accumulation of pS129 in various SNCA triplication models. Oliveira et al. (2015) observed elevated α-synuclein in dopaminergic neurons derived from 3xSNCA iPSCs, while Jo et al. (2021) identified Lewy body–like inclusions in 3xSNCA midbrain organoids at day 90 following treatment with the lysosomal inhibitor conduritol B epoxide (CBE). Mohamed et al. (2021) reported increased aggregation of oligomeric and phosphorylated α-synuclein in 3xSNCA midbrain organoids by day 100, with pathological progression through day 170. Becerra-Calixto et al. (2023) showed progressive pS129 accumulation in 3xSNCA midbrain organoids from days 120 to 180, while Muwanigwa et al. (2024) detected pS129-rich aggregates in 3xSNCA human midbrain organoids by day 70.

While these studies describe accumulation and progressive pathology markers, our 3xSNCA assembloid model demonstrates robust pS129-positive aggregate formation with pathological morphology, as defined by Lashuel et al. (2022), by day 50. This significantly accelerated aggregation highlights the critical role of 3xSNCA microglia in promoting pathological α- synuclein accumulation. This early pathology is likely driven by an overproduction of α-synuclein that overwhelms cellular clearance systems such as chaperone-mediated autophagy, the ubiquitin–proteasome system, and macroautophagy (Cuervo et al., 2004; Martinez-Vicente et al., 2008; Snyder et al., 2003; Winslow et al., 2010), resulting in accumulation of misfolded α- synuclein and aggregate formation.

Importantly, 3xSNCA microglia induced pS129 pathology even when integrated into wild-type midbrain organoids (chimeric assembloids), demonstrating that microglial dysfunction alone can initiate α-synuclein pathology independent of neuronal SNCA overexpression. These finding positions microglia as active drivers of disease initiation and progression, rather than passive responders (Zhang et al., 2005; Hickman et al., 2018; Ferreira & Romero-Ramos, 2018; Bido et al., 2021).

Given the prominent α-synuclein pathology observed in the here described model, these findings offer a valuable platform to explore therapeutic strategies that directly target α-synuclein aggregation or modulate downstream disruptions in proteostasis. Several small molecules have shown promising results in this context. Squalamine and Trodusquemine displace α-synuclein from lipid membranes, effectively blocking both the initiation and propagation of aggregation in preclinical models (Perni et al., 2017; Rao et al., 2000; Yang et al., 2024). Minzasolmin (UCB0599) prevents α-synuclein misfolding, while Emrusolmin (Anle138b) binds to oligomeric forms to inhibit the formation of toxic aggregates. Both compounds have demonstrated efficacy in preclinical studies and are undergoing clinical evaluation (Price et al., 2023; Heras-Garvin et al., 2019; Wegrzynowicz, et al., 2019; Levin et al., 2022). In parallel, strategies that enhance lysosomal degradation of α-synuclein may offer complementary benefits. Ambroxol, for example, boosts glucocerebrosidase activity and lysosomal function, promoting clearance of misfolded protein and reducing oxidative stress (Sechi & Sechi, 2024). Understanding how pathological processes are shaped by microglia is particularly important, as these cells can act as both contributors to and modulators of neurodegeneration. To unravel their complex role, there is a critical need for physiologically relevant *in vitro* models that capture the most essential functional properties of microglia. These therapeutic candidates, with their distinct mechanisms of action, offer promising avenues to intervene in disease-relevant pathways.

Taken together, our findings highlight that the 3xSNCA assembloid model recapitulates key features of Parkinson’s disease, including spontaneous pS129 accumulation and microglia-driven α-synuclein pathology. By modelling microglia–neuron interactions without exogenous fibrils, the assembloid and chimeric systems offer physiologically relevant platforms to investigate early drivers of synucleinopathies and evaluate therapeutic strategies.

### Data availability

All original and processed data, along with the scripts supporting the findings of this study, are publicly available at this https://doi.org/10.17881/syhp-k282. Gene expression datasets have been deposited in the Gene Expression Omnibus (GEO) under the accession number GSE299260.

### Code availability

All scripts used to obtain, analyse and plot the data are available at https://gitlab.com/uniluxembourg/lcsb/developmental-and-cellular-biology/zuccoli_2025.

## Acknowledgments

Image and flow cytometry data acquisition were supported by the LCSB Bioimaging Platform (BIP, RRID:SCR_026352), including its flow cytometry core, at LCSB (RRID:SCR_026168) of the University of Luxembourg (RRID:SCR_011632). We acknowledge Novogene for their RNA sequencing service and we would also like to thank Daniela Frangenberg-Hoff, Raquel Forte Marques and Mona Tuzza for excellent technical assistance.

This project has received funding from the i2TRON Doctoral Training Unit (PRIDE19/14254520) which is funded by the Luxembourg National Research Fund (FNR).

Rights retention statement: This research was funded in whole by the FNR-Luxembourg. For the purpose of Open Access, the author has applied a CC BY public copyright license to any Author Accepted Manuscript (AAM) version arising from this submission.

## Author Contributions

E.Z. conceived, designed, collected data, performed data analysis, and interpretation of results. S.S.S., A.Z. and I.R. performed data analysis. H.K., A.S.Z. contributed with experiments. The work was supervised by J.C.S., E.Z. wrote the original manuscript. E.Z., H.K., S.S.S., A.Z., I.R., A.S.Z and J.C.S. revised and edited the manuscript.

---

**Supplementary Figure 1: iPSC-derived microglia express specific microglial markers.** (a) Representative immunofluorescence staining of mature (day 14) WT cell lines (WT1, WT2 and WT3), 3xSNCA and KO microglia for macrophage markers TMEM119 (Transmembrane Protein 119) and CD45, (b) the macrophage marker PU.1, (c) the microglial marker IBA1 as well as Hoechst 33342 (scale bar 20 μm, 63x).

**Supplementary Figure 2: α-synuclein expression in WT lines and transcriptomic alterations in 3xSNCA microglia.** (a) Representative immunofluorescence staining of additional WT cell lines showing the microglial marker P2RY12 (Purinergic Receptor P2Y12), total α-synuclein and nuclei (Hoechst 33342) (scale bar 20 μm, 63x). (b) Volcano plot showing differential gene expression (DEGs) between 3xSNCA and WT microglia. Genes with log_2_FC>2 and p.adjust <0.01 are highlighted. (c) Unsupervised hierarchical clustering of 3xSNCA and WT samples based on normalised gene counts of top significant DEGs involved in the inflammasome pathway.

**Supplementary Figure 3: Co-localization of aggregation markers with α-synuclein and pS129 in 3xSNCA ASBs.** (a) Representative image of 3xSNCA ASBs to show pS129 antibody specificity. Sections were stained with pS129 [MJF-R13 (8-8)] and Hoechst 33342 (scale bar 100 μm, 20x; 20 μm, 63x; 5 μm, zoom in of inset). (b) Representative image of WT and 3xSNCA ASBs stained with total α-synuclein, pS129 [MJFR-14-6-4-2)] and the aggregation marker Thioflavin S (scale bar 100 μm, 20x; 20 μm, 63x). (c) Zoom in of inset of representative image and 3D reconstruction of 3xSNCA ASBs stained with pS129 [MJFR-14-6-4-2)] and the aggregation marker Thioflavin S (5 μm, zoom in of inset). (d) High-content automated image analysis of immunofluorescence stainings showing the foldchange of p62+ cells and (e) Proteostat+ cells normalized by total nuclei in WT and 3xSNCA ASBs. Data represents three independent batches (n=3) normalized to the mean of the controls per batch. Mann-Whitney *U* test; **p < 0.01, ***p < 0.001. (f) Representative image of WT and 3xSNCA ASBs stained with total α-synuclein, the mitochondrial marker TOM20 and Hoechst 33342 (scale bar 100 μm, 20x; 20 μm, 63x). (g) *Left panel:* The same representative image of 3xSNCA ASB as shown in (f), displayed here with total α-synuclein and TOM20 markers (Hoechst omitted). *Right panel:* 3D reconstruction of this image using IMARIS software (scale bar, 5 μm; zoom-in of inset).

**Supplementary Figure 4: WT-3xSNCA ASBs show co-localization of aggregation markers with α-synuclein and pS129 antibodies.** (a) High-content automated image analysis of immunofluorescence stainings showing the foldchange of p62+ cells and (b) Proteostat+ cells normalized by total nuclei in WT and WT-3xSNCA ASBs. Data represents three independent batches (n=3) normalized to the mean of the controls per batch. Mann-Whitney *U* test; **p < 0.01. (c) Representative image of WT and WT-3xSNCA ASBs stained with pS129, Ubiquitin and Hoechst 33342 (scale bar 100 μm, 20x; 20 μm, 63x). (d) *Left panel:* The same representative image of WT-3xSNCA ASB as shown in (c), displayed here with pS129 and Ubiquitin markers (Hoechst omitted). *Right panel:* 3D reconstruction of this image using IMARIS software (scale bar, 5 μm; zoom-in of inset). (e) Representative image of WT and WT-3xSNCA ASBs stained with total α-synuclein, the mitochondrial marker TOM20 and Hoechst 33342 (scale bar 100 μm, 20x; 20 μm, 63x). (f) *Left panel:* The same representative image of WT-3xSNCA ASB as shown in (e), displayed here with total α-synuclein and TOM20 markers (Hoechst omitted). *Right panel:* 3D reconstruction of this image using IMARIS software (scale bar, 5 μm; zoom-in of inset).

