## Supplementary figures and images for "Parkinson’s disease microglia induce endogenous α-synuclein pathology in patient-specific midbrain organoids"

### Supplementary Figures

#### Supplementary Figure 1

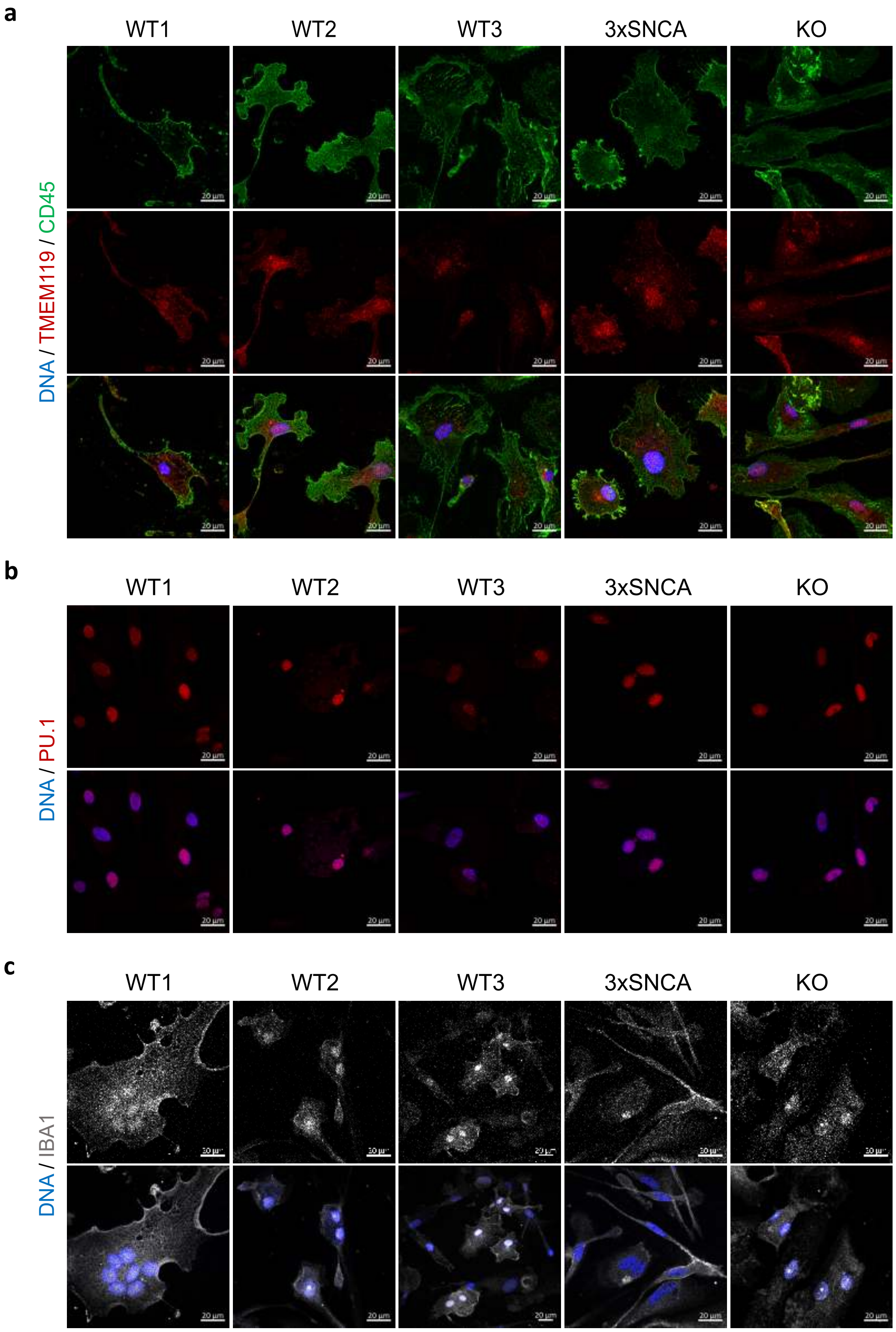

**Supplementary Figure 2**

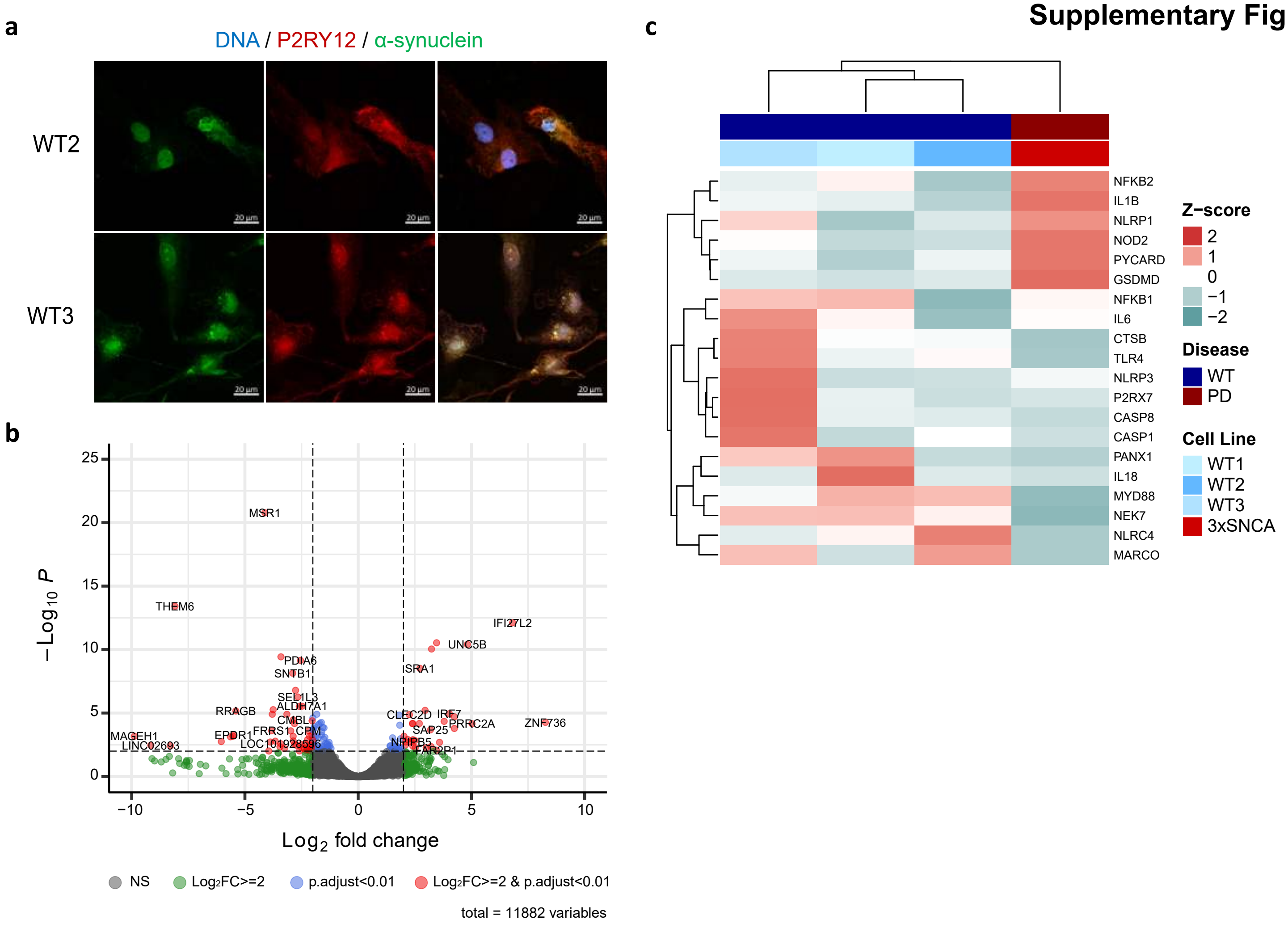

**Supplementary Figure 3**

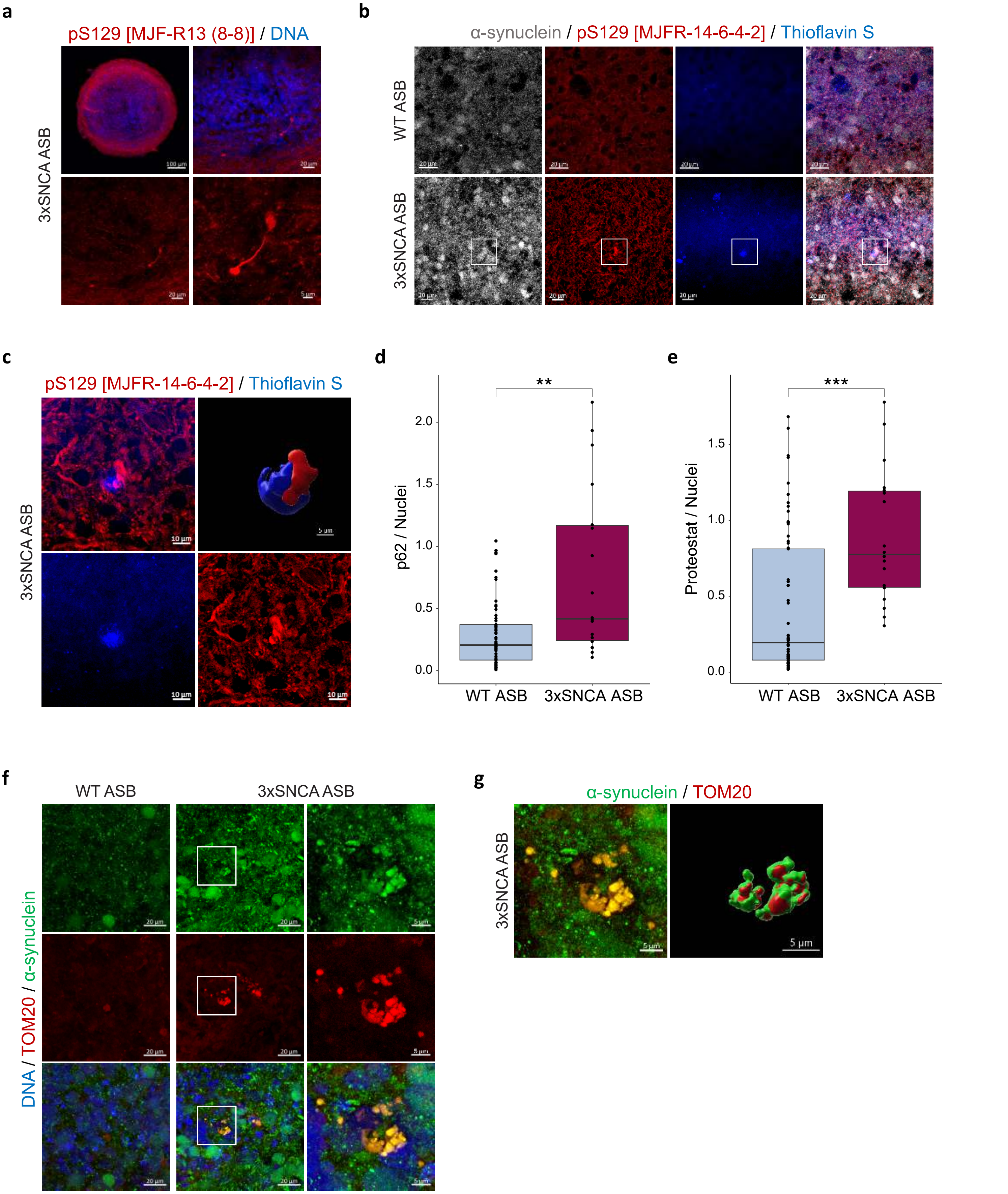

**Supplementary Figure 4**

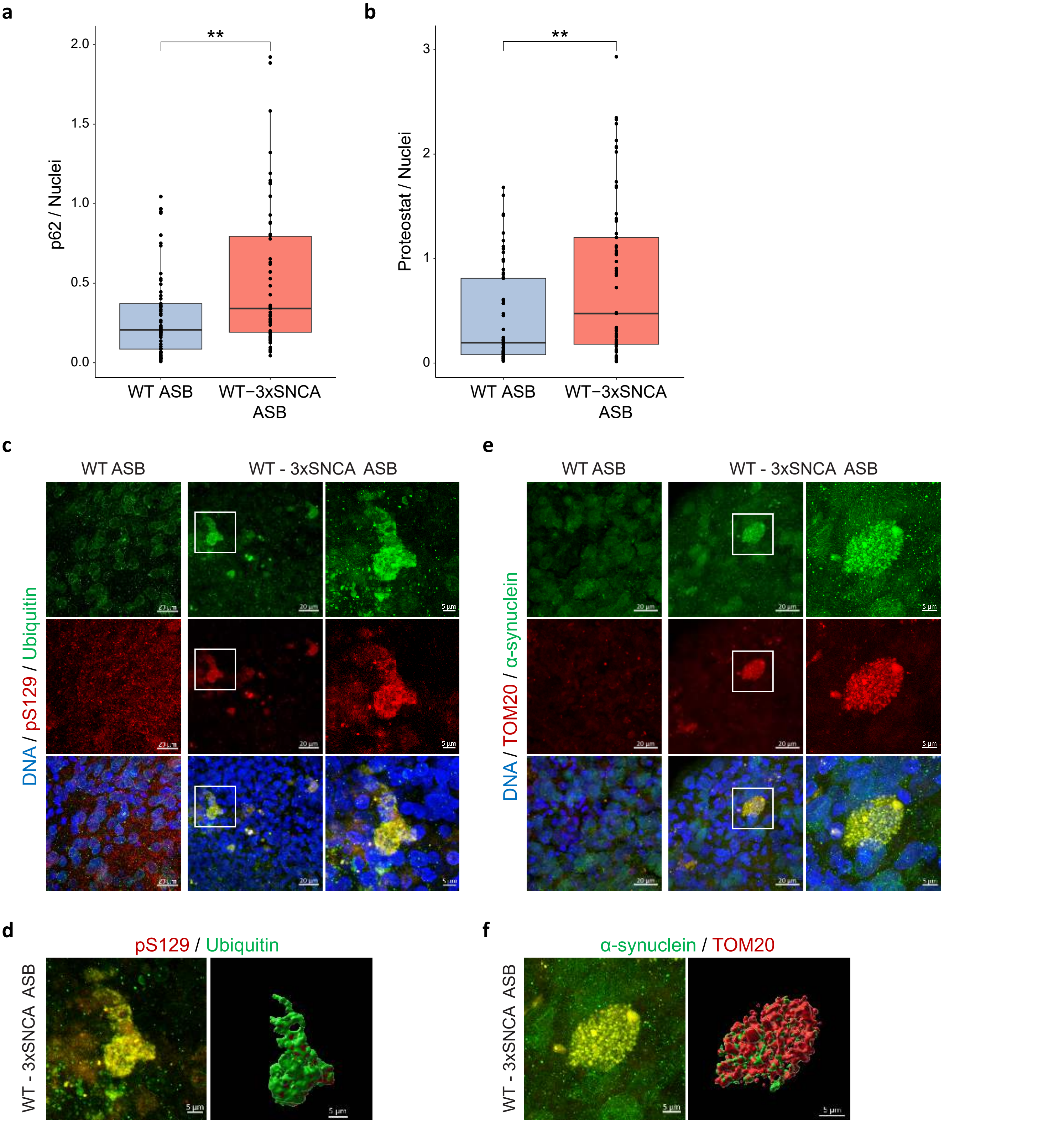
