## Supplementary Table 1 for "Parkinson’s disease microglia induce endogenous α-synuclein pathology in patient-specific midbrain organoids"

| Lab Identifier | Sample ID | Diagnosis | Patient | Sex | Age of sampling | Source of iPSC |
| --- | --- | --- | --- | --- | --- | --- |
| 201 | WT1 | Healthy | A13777 | F | - | Gibco |
| 232 | WT2 | Healthy | T12.9/C1-2 | F | 53 | Reinhardt et al. 2013 |
| 277 | WT3 | Healthy | 2716623-MDPD1-CHL | F | 65 | IBBL / Muenster |
| 336 | 3xSNCA | PD | ND27760 | F | 55 | European Bank for induced pluripotent stem cells - EDi001-A |
| 374 | KO | PD | ND27760 | F | 55 | Chen et al. 2019 |

**Supplementary Table 1. Cell lines used in this study**
