## Supplementary Table 2 for "Parkinson’s disease microglia induce endogenous α-synuclein pathology in patient-specific midbrain organoids"

**Supplementary Table 2: Primary and secondary antibodies used in western blot and dot blot.**

| Antibody | Source | Cat. no. | RRID | Species | Dilution | Method |
| --- | --- | --- | --- | --- | --- | --- |
| β-Actin | Cell Signaling Technology | 3700S | *AB_2242334* | mouse | 1:20000 | WB |
| α-synuclein (2A7) | Novus Biologicals | NBP1-05194 | *AB_1555287* | mouse | 1:1000 | WB |
| α-synuclein [MJFR1] | Abcam | ab138501 | *AB_2537217* | rabbit | 1:1000 | Dot Blot |
| pS129 (D1R1R) | Cell Signaling Technology | 23706S | *AB_2798868* | rabbit | 1:1000 | WB |
