## Supplementary Table 3 for "Parkinson’s disease microglia induce endogenous α-synuclein pathology in patient-specific midbrain organoids"

**Supplementary Table 3: Primary and secondary antibodies used in immunofluorescence stainings.**

| Antibody | Source | Cat. no. | RRID | Species | Dilution |
| --- | --- | --- | --- | --- | --- |
| TMEM119 | Sigma-Aldrich | HPA051870 | *AB_2681645* | rabbit | 1:250 |
| CD45 | BioLegend | 304002 | *AB_314390* | mouse | 1:1000 |
| PU.1 (9G7) | Cell Signaling Technology | 2258 | *AB_2186909* | rabbit | 1:250 |
| IBA1 | Abcam | ab5076 | *AB_2224402* | goat | 1:250 |
| P2RY12 | Sigma-Aldrich | HPA014518 | *AB_2669027* | rabbit | 1:250 |
| MAP2 | Abcam | ab92434 | *AB_2138147* | chicken | 1:1000 |
| TH | Abcam | ab112 | *AB_297840* | rabbit | 1:1000 |
| S100β | Sigma-Aldrich | S2532 | *AB_477499* | mouse | 1:600 |
| GFAP | Millipore | MAB3402 | *AB_94844* | mouse | 1:1000 |
| α-synuclein (2A7) | Novus Biologicals | NBP1-05194 | *AB_1555287* | mouse | 1:1000 |
| α-synuclein [MJFR1] | Abcam | ab138501 | *AB_2537217* | rabbit | 1:1000 |
| pS129 (D1R1R) | Cell Signaling Technology | 23706S | *AB_2798868* | rabbit | 1:1000 |
| pS129 [MJF-R13 (8-8)] | Abcam | ab168381 | *AB_2728613* | rabbit | 1:500 |
| pS129 [MJFR-14-6-4-2] | Abcam | ab209538 | *AB_2714215* | rabbit | 1:500 |
| SQSTM1 / p62 | Abcam | ab155686 | *AB_2847961* | rabbit | 1:500 |
| Ubiquitin (P4D1) | Santa Cruz Biotechnology | sc-8017 | *AB_628423* | mouse | 1:500 |
| Tom20 (D8T4N) | Cell Signaling Technology | 42406 | *AB_2687663* | rabbit | 1:300 |
| Anti-mouse 488 | Invitrogen | A32766 | *AB_2762823* | Donkey | 1:1000 |
| Anti-mouse 647 | Invitrogen | A-31571 | *AB_162542* | Donkey | 1:1000 |
| Anti-rabbit 488 | Invitrogen | A21206 | *AB_2535792* | Donkey | 1:1000 |
| Anti-rabbit 568 | Invitrogen | A-10042 | *AB_2534017* | Donkey | 1:1000 |
| Anti-chicken 488 | Jackson Immunoresearch | 703-545-155 | *AB_2340375* | Donkey | 1:1000 |
| Anti-goat 647 | Invitrogen | A32849 | *AB_2762840* | Donkey | 1:1000 |
